# Cftr Modulates Wnt/β-Catenin Signaling and Stem Cell Proliferation in Murine Intestine

**DOI:** 10.1101/156562

**Authors:** Ashlee M. Strubberg, Jinghua Liu, Nancy M. Walker, Casey D. Stefanski, R. John MacLeod, Scott T. Magness, Lane L. Clarke

## Abstract

**Background & Aims:** Cystic fibrosis (CF) patients and CF mouse models have increased risk for gastrointestinal tumors. CF mice exhibit augmented intestinal proliferation of unknown etiology and an altered intestinal environment. We examined the role of Cftr in Wnt/β-catenin signaling, stem cell proliferation and its functional expression in the active intestinal stem cell (ISC) population. Dysregulation of intracellular pH (pH_i_) in CF ISCs was investigated for facilitation of Wnt/β-catenin signaling.

**Methods:** Crypt epithelia from wild-type (WT) and CF mice were compared ex vivo and in intestinal organoids (enteroids) for proliferation and Wnt/β-catenin signaling by standard assays. Cftr in ISCs was assessed by immunoblot of sorted Sox9^EGFP^ intestinal epithelia and pH_i_ regulation by confocal microfluorimetry of Lgr5+-EGFP ISCs. Plasma membrane association of the Wnt transducer Disheveled 2 (Dvl2) was assessed by fluorescence imaging of live enteroids from WT and CF mice crossed with Dvl2-EGFP/Rosa^mT/mG^ mice.

**Results:** Relative to WT, CF intestinal crypts showed a ~30% increase in epithelial and Lgr5+ ISC proliferation and increased Wnt/β-catenin signaling. Cftr was expressed in Sox9^EGFPLo^ ISCs and loss of Cftr induced an alkaline pH_i_ in Lgr5+-EGFP ISCs. CF crypt-base columnar cells (CBCs) demonstrated a generalized increase in plasma membrane Dvl2-EGFP association as compared to WT. Dvl2-EGFP membrane association was charge- and pH-dependent and increased in WT CBCs by Cftr inhibition.

**Conclusions:** CF intestine exhibits increased ISC proliferation and Wnt/β-catenin signaling. Loss of Cftr increases pH_i_ in ISCs which stabilizes the plasma membrane association of the Wnt transducer Dvl, likely facilitating Wnt/β-catenin signaling. Absence of Cftr-dependent suppression of ISC proliferation in the CF intestine may contribute to increased risk for intestinal tumors.

Graphical Abstract

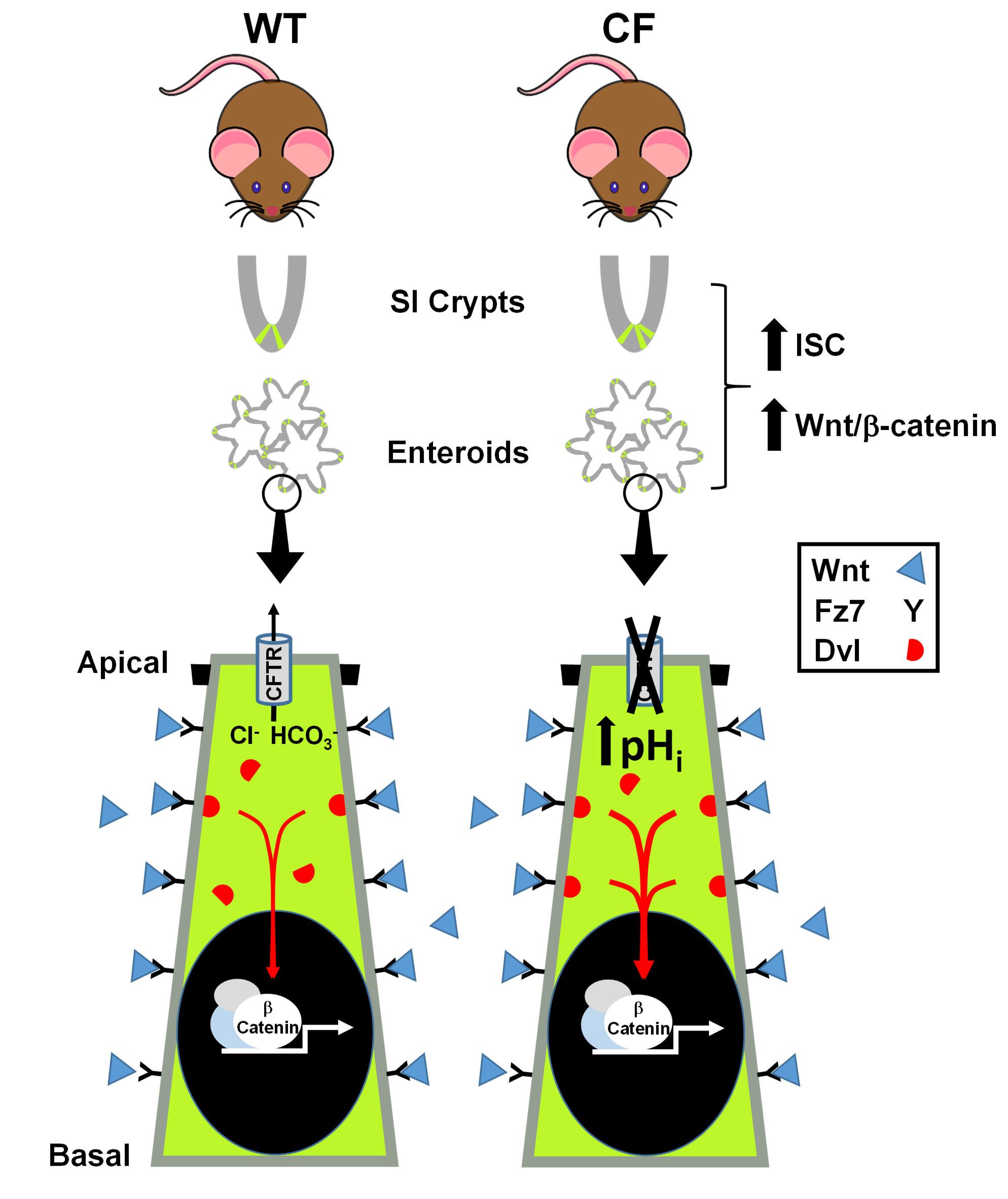

## Introduction

Cystic fibrosis (CF) is a heritable genetic disease caused by mutations in the cystic fibrosis transmembrane conductance regulator gene. The gene product (CFTR) is a major anion channel of fluid transporting epithelia where it functions in transepithelial Cl^−^ and HCO_3_- secretion^1, 2^. CF affects multiple organs, in particular the airway epithelia where failure of mucociliary clearance results in bacterial colonization of the lung.

However, intestinal disease is one of the earliest manifestations of CF and presents life-long conditions, including small intestinal bacterial overgrowth (SIBO)^3^, low-grade small bowel inflammation^4, 5^, obstructive bowel disease^6, 7^ and an increased incidence of gastrointestinal (GI) cancer^8, 9^. Cftr knockout (KO) mice recapitulate CF intestinal disease without significant manifestations of pancreatic, liver or lung disease. The disease phenotype includes a high incidence of bowel obstruction^10^, low-grade bowel inflammation^11^, small intestinal bacterial overgrowth^3^, dysbiosis^12^ and the spontaneous development of intestinal tumors with age, inclusive of invasive forms^13^.

Previous in vivo studies show that Cftr KO mice exhibit increased intestinal epithelial proliferation without a corresponding increase in apoptosis^14^, a condition that may predispose to intestinal neoplasia^15^. Recent epidemiological studies reveal strong correlations between the rate of stem cell division and the incidence of cancer^16^. Since the intestine has one of the highest rates of epithelial turnover in the body, pathological manifestations of CF that enhance the rate of epithelial turnover and contribute to intestinal inflammation are predicted to increase the risk of gastrointestinal cancer.

However, a mechanistic understanding linking the absence of Cftr with enhanced proliferation of the intestinal epithelium, particularly the stem cell population, has not been advanced.

Cftr is highly expressed in intestinal crypts^17, 18^, the proliferative compartment of the intestine, and by providing apical membrane Cl^−^ and HCO_3_- ion permeability, has an impact on the regulation of epithelial intracellular pH (pH_i_). Loss of Cftr function by acute channel blockade or in Cftr KO enteroids results in an incompletely compensated alkaline pH_i_ in the crypt epithelium19. Compensation of the alkaline pH_i_ is impaired by a corresponding increase in intracellular Cl^−^ concentration [Cl^−^]_i_, which reduces cellular anion exchange activity^20^. Several aspects of cell proliferation are known to be facilitated by an alkaline pH_i_, including cell cycle phase progression at G2/M^21^, optimization of DNA replication^22^, cytoskeleton remodeling and cell migration^23,24^ and membrane biogenesis^25^. Cell alkalinity has also been shown to facilitate Wnt signaling^26, 27^, which may directly affect stem cell proliferation.

Wnt/β-catenin signaling is essential for homeostasis and proliferation of intestinal stem cells28 and is often aberrantly activated in intestinal cancer. In *Drosophila* spp., pH_i_ changes can alter Wnt signaling by modulating the interaction of the initial signal mediator Disheveled (Dvl) with the Wnt receptor Frizzled (Fz) at the plasma membrane^26^. The critical binding of Dvl’s PDZ domain with the PDZ binding domain of Fz is facilitated by a stable interaction of Dvl’s polybasic DEP domain to negatively charged phospholipids (phosphatidic acid, phosphatidylglycerol) at the inner leaflet of the plasma membrane. Phospholipid interaction is pH_i_- and charge-dependent such that proton electrostatic interference at an acidic pH_i_ reduces DEP domain membrane binding and subsequent Wnt signaling26. We hypothesized that an alkaline pH_i_ in Cftr KO intestinal stem cells (ISCs) stabilizes Dvl interaction at plasma membrane thereby facilitating Wnt/β-catenin signaling.

The present study investigates augmented proliferation of the intestinal epithelium in a Cftr KO mouse model. Studies first examine whether hyperproliferation persists in Cftr KO enteroid culture, which isolates the epithelium from the immediate consequences of an abnormal Cftr KO intestinal environment (inflammation, dysbiosis) and provides the technological advantage of live crypt cell imaging^3, 11, 29^. Second, studies evaluate the activation status of Wnt/β-catenin signaling in the Cftr KO intestine and the functional activity of Cftr in ISCs, specifically, Lgr5+ stem cells^30^. Third, live cell imaging is used to examine the hypothesis that alkalinity of Cftr KO intestinal crypt base columnar stem cells is conducive to increased interaction of Dvl (i.e., the major isoform Dvl2^31^) with the plasma membrane for Wnt signaling.

## Materials and Methods

### Mice

Mice with gene targeted disruptions of the murine homolog of *Cftr* [*abcc7*, Cftr knock-out (KO)] and sex-matched wild-type (WT, +/+ or +/-) littermates were used^10^. Mice were outbred to Black Swiss (Charles River) mice at generational intervals and resultant F1 heterozygotes were crossed to generate F2 offspring for experimentation. The Cftr KO mouse line was crossed with Lgr5-EGFP-IRES-creERT2 (Lgr5-EGFP, Jackson Labs) mice to generate WT/Lgr5-EGFP and Cftr KO/Lgr5-EGFP mice. The Cftr KO mouse line was also crossed with both Dvl2 KO/Dvl2-EGFP BAC transgenic^32^ and Gt(ROSA)^26Sortm4(ACTB-tdTomato,-EGFP)Luo^/J (Rosa^mT/mG^, Jackson Labs) mouse lines to generate WT and Cftr KO/Dvl2 KO/Dvl2-EGFP/Rosa^mT/mG^ mice. Genotypes were identified by PCR analysis of tail-snip DNA as previously described for mutant Cftr^33^, Dvl2 KO and Dvl2-EGFP expression^34^, and Rosa^mT/mG^ (Jackson Labs). Copy number for the Dvl2-EGFP transgene was verified by TaqMan^®^ GFP copy number assay (ThermoFisher Scientific). Only mice expressing 2 copies of the Dvl2-EGFP transgene were used for experimental analysis. All mice were maintained *ad libitum* on standard laboratory chow (Formulab 5008, Rodent Chow; Ralston Purina) and distilled water containing Colyte^®^ (Schwartz Pharma) laxative to prevent intestinal obstruction in the Cftr KO mice. Mice were housed individually in a temperature- and light-controlled room (22–26°C; 12-hour light: 12-hour dark cycle) in the Association for Assessment and Accreditation of Laboratory Animal Care-accredited animal facility at the Dalton Cardiovascular Research Center, University of Missouri. Mouse experiments were performed in accordance with guidelines outlined in “Guide for the Care and Use of Laboratory Animals” prepared by the National Academy of Sciences and published by the National Institutes of Health and with approval from the University of Missouri IACUC.

### Enteroid culture

The enteroid culture of isolated crypt epithelium from the proximal jejunum has been previously described in detail^19^. Cultures were overlaid with growth medium containing Ham’s F-12 medium with 5% FBS, 50µg/mL gentamicin, 125ng/mL R-spondin1, 25ng/mL noggin and 12.5ng/mL epidermal growth factor. Growth medium was changed every 3–4 days and enteroids were passaged every 7–10 days using Cell Recovery Solution (BD Sciences). Except where otherwise indicated, passages 1–2 were used for experimentation.

### Lgr5-EGFP cell counts

Crypts were isolated from WT/Lgr5-EGFP or Cftr KO/Lgr5-EGFP mice and fixed immediately in 10% buffered formalin (Sigma-Aldrich), or cultured as enteroids in growth medium before fixation. Z-stack images were acquired using either a TCS SP5 Confocal-Multiphoton microscope built on a DMI6000 inverted platform (Leica, Wetzler, Germany) or an Olympus Fluoview confocal microscope (FV1000). Images were reconstructed in 3D using Imaris^®^ Software (version 7.7.1, Bitplane, Concord, MA) and the number of Lgr5-EGFP positive stem cells identified in three-dimensions was counted for each crypt.

### Immunofluorescence

Freshly isolated crypts or enteroids from the proximal jejunum were fixed in 10% buffered formalin (Sigma-Aldrich) or 4% paraformaldehyde (PFA) and stored at 4°C until processing. Fixed enteroids and Matrigel™ were scraped from the culture dishes, transferred to 1.5mL tubes, centrifuged at 200*g* (1min) and washed 3x with 1X PBS to remove Matrigel™ and fixative. Samples were permeabilized for 60mins using 0.5% Triton X-100 (Sigma-Aldrich) in PBS and blocked for 30min with gentle shaking in fish skin gelatin buffer [10mM Tris, 5mM EDTA, 0.15M NaCl, 0.25% fish skin gelatin (Sigma-Aldrich), 0.05% Tween20]. Samples were incubated overnight at 4°C with primary antibody diluted in fish skin gelatin buffer, washed 3x in fish skin gelatin buffer, and incubated with secondary antibody for 4–5 hours with gentle shaking at 4°C. After removal of the secondary antibody, samples were washed three times for 10mins in fish skin gelatin buffer. Samples were then resuspended in SlowFade® gold antifade mounting medium (ThermoFisher Scientific) and sealed under glass coverslips on microscope slides (ThermoFisher Scientific). Fresh crypts and enteroids were imaged using an Olympus Fluoview confocal microscope. Z-stacks were imaged to determine crypt cross-sections post-acquisition using Imaris® software. Anti-Frizzled 7 (10µg/mL, R&D Systems, AF198) was used as primary antibody and secondary antibody was anti-goat IgG Alexa Fluor® 405 (Abcam, ab175665) used at a 1:500 dilution in fish skin gelatin buffer.

### Proliferation assays

Proliferation of WT and Cftr KO freshly isolated crypts was measured by immunofluorescence for mitotic cells using anti-Phospho-Histone H3 (PH3) primary antibody (1:100 dilution, Millipore, 06-570) in fish skin gelatin buffer (see methods above). Nuclei were labeled with TO-PRO 3 (ThermoFisher Scientific) nuclear stain (1:2000 dilution) in fish skin gelatin buffer. Proliferation of enteroid crypts (p1 or p2, 5–7 days) maintained in growth medium was measured using EdU to label cells in S phase of the cell cycle. Enteroids were exposed in situ to EdU for 15mins, fixed in 4% PFA and stored at 4°C. Fixed enteroids and Matrigel™ were scraped from the culture dishes, transferred into 1.5mL tubes, and centrifuged at 200*g* for 1min. The supernatant containing Matrigel™ was aspirated and the fixed enteroids were washed twice with 1X PBS. The Click-iT 5-ethynyl-2’-deoxyuridine (EdU) assay was performed according to the manufacturer’s protocol (ThermoFisher Scientific) and as previously described^35^. Nuclei were labeled with Hoechst 33342 diluted 1:2000 for 1h. Labeled enteroids were concentrated by brief centrifugation (200*g*, 1min), resuspended in SlowFade® gold antifade mounting medium (ThermoFisher Scientific) and sealed under a glass coverslip on microscope slides. Crypts were imaged on an Olympus Fluoview confocal microscope (FV1000). Post-acquisition 3D reconstructed Z-stacks (Imaris® Software, Bitplane) were used to determine crypt cross sections for counting of PH3-positive (PH3+), EdU-positive (EdU+) and total nuclei. Acquired images were coded and nuclei counting was performed by an observer blinded to genotype.

### Immunoblot analysis

Freshly isolated crypts, enteroids, or sorted intestinal epithelial cells were suspended in ice-cold, radioimmunoprecipitation assay (RIPA) buffer (Cell Signaling) containing Halt™ Protease inhibitor (Thermo Fisher Scientific) and lysed at 4°C by supersonication. Total lysate protein was loaded on 10% SDS-PAGE gels for electrophoresis, membrane transfer and immunoblotting. Anti-active β-catenin (1:2000 dilution, Millipore, 05-665), anti-Lef1 (1:500 dilution, Santa Cruz, sc-28687) and anti-Cftr 3G11 (1:2000 dilution, provided by CFTR Folding Consortium, Cystic Fibrosis Foundation Therapeutics) were used as primary antibodies. Anti-β-actin (1:2000 dilution, Santa Cruz, sc-130656) or anti-GAPDH (1:2000 dilution, Santa Cruz, sc-25778) were used as loading controls. Densitometry was performed using Image Lab™ Software (version 5.2.1, BioRad).

### FACS

Isolated small intestinal crypt cells from Sox9^EGFP^ mice were dissociated for fluorescence-activated cell sorting, as previously described^36^. Sox9EGFP^Negative^, Sox9^EGFPSublo^, Sox9^EGFPLo^ and Sox9^EGFPHigh^ cell were isolated using a MoFlo FACS machine (Dako/Cytomation). Cell were collected in RIPA buffer and frozen at −20°C for protein immunoblotting.

### Confocal microfluorimetry of intracellular pH

WT, Cftr KO, WT/Lgr5-EGFP and Cftr KO/Lgr5-EGFP enteroids were cultured in growth medium for 5–7 days on glass-bottomed Fluorodishes (World Precision Instruments). Enteroids were loaded with the ratiometric pH sensitive dye SNARF-5F (Thermo Fisher Scientific) at 40µM for 30min at 37°C as previously described^19^. For basal conditions, cultures were continuously superfused with Kreb’s bicarbonate Ringer (KBR) + 5mM N-tris(hydroxymethyl)-methyl-2-aminoethanesulfonic acid (TES) buffer and gassed with 95% O_2_: 5% CO_2_ (pH 7.4, 37°C). For Cftr KO enteroids grown in pH 7.1 or 6.6 medium, enteroids were imaged in static culture medium (pH 7.1 or 6.6) with an overlying 95% air: 5% CO_2_ atmosphere maintained at 37°C using a culture dish incubator DH-35iL (Warner Instruments, 64-0349). All images were acquired with a TCS SP5 Leica Confocal microscope in a temperature controlled incubator. The excitation source for SNARF-5F was a 514 nm argon laser and images were collected at dual emission wavelengths (580 ± 30 and 640 ± 30 nm). Z-stacks of individual crypts were imaged and regions of interest (ROI) were placed post-acquisition using Imaris® Software. For intracellular regional pH_i_ studies, ROI were placed on cross-sectional slices at the apical aspect of CBCs near the +4 position (to avoid Paneth cell granule fluorescence) using SlideBook 5.0 (Intelligent Imaging Innovations, Denver, CO). The 580-to-640-nm ratio was converted to pH_i_ using a standard curve generated by the K^+^/nigericin technique in unlabeled and Lgr5-EGFP-positive cells^37^.

### Live imaging of enteroids for Dvl2-EGFP membrane association

Passaged enteroids were plated in Matrigel™ onto Fluorodishes and cultured for 5–7 days prior to imaging. Unless indicated otherwise, enteroids were gassed with 95% air: 5% CO_2_ and maintained at 37°C using a culture incubator DH-35iL during all acquisitions. Z-stacks were acquired using an Olympus Fluoview 1000 confocal microscope. Post-acquisition 3D reconstructed Z-stacks (Imaris® Software, Bitplane) were used to determine crypt cross sections for Dvl2-EGFP proximity to the plasma membrane of CBCs in live enteroid crypts. CBCs were selected based on 1) location at cell positions 1–3; 2) lack of granulation as defined by the <u>absence</u> of a granule theca outlined by Rosa^mT/mG^ label (prominent in Paneth or goblet cells) and lack of granules by light images; 3) a contiguous and well-defined Rosa^mT/mG^ labeled plasma membrane; and 4) not overtly undergoing cell division as indicated by a nuclear position apical to the basal membrane and membrane accumulation at the apical pole of the cell. Using ImageJ software (version 1.49, Bethesda, MD), preliminary studies indicated that Dvl2-EGFP intensity measurements taken at CBC plasma membrane adjacent to the nucleus were confounded technically (due to minimal cytoplasm and curvature of the nucleus) and conceptually (apposition of nuclear and plasma membranes). Therefore, measurement of the juxtamembrane Dvl2-EGF intensity was confined to the lateral membranes in the supranuclear portion of CBCs, i.e., apical to the nucleus. Acquired images were coded and measurement of juxtamembrane Dvl2-EGFP was performed by an observer blinded to genotype. To quantitate membrane proximity of Dvl2-EGFP at the supranuclear lateral plasma membranes of CBC, the Rosa^mT/mG^ labeled lateral plasma membranes were outlined as a ROI on the red (mTomato) channel images of cell cross-sections using ImageJ software. The ROI was copied to the green (EGFP) channel image of the CBC for measurement of Dvl2-EGFP intensity within a two-pixel distance (1.15µm) interior to the supranuclear lateral plasma membrane, and avoiding the apical (brush border) membrane. The average Dvl2-EGFP intensity of pixels was also acquired for the entire supranuclear region of each CBC. This measurement was used to normalize differences in EGFP intensity between CBCs by calculating the ratio of average juxtamembrane EGFP intensity divided by average EGFP intensity of all pixels in the supranuclear region of the CBC. Average EGFP intensity of pixels in the supranuclear region were not significantly different between WT and Cftr KO CBCs (Ave. intensity: WT = 59.5±8.8; and Cftr KO = 88.5±15.9, ns, n = 7–6, respectively). Because measurements of juxtamembrane apposition of Dvl2-EGFP were performed on 2D confocal slices of CBCs, EGFP intensity at the lateral membrane oriented lateral (lateral-Lateral) and medial (medial-Lateral) to the central axis of the crypt were also individually collected (see Fig. 4C, 4D).

### Charge- and pH_i_-dependence of Dvl2-EGFP association with the plasma membrane in Cftr KO enteroids

Cftr KO enteroids were treated with either sphingosine (Avanti Polar Lipids), low bicarbonate medium, bumetanide plus carbachol (CCH, Sigma) or Cftr_inh_-172 (Sigma) plus GlyH-101 (Millipore) to alter Dvl2-EGFP localization at the plasma membrane. For experiments using sphingosine to neutralize negative charges at the inner leaflet of the plasma membrane, Cftr KO enteroids were treated with sphingosine in growth medium for 1hr prior to imaging (95% ethanol: 5% water used as vehicle, 1:2000 final dilution). For experiments lowering pH_i_, Cftr KO enteroids were cultured for 3 days in standard medium and then 2 days in either in low bicarbonate (3.6mM bicarbonate, 7mM sodium TES, 6.8mM mannitol, pH 6.6) medium or maintained in standard medium (14mM bicarbonate, pH 7.1). For experiments to acutely reduce intracellular Cl^−^ and pH_i_, Cftr KO enteroids were sequentially treated with bumetanide (50µM, Sigma; dissolved in 95% ethanol) and CCH (100µM), as previously described^20^. Briefly, Cftr KO enteroids were treated with bumetanide for 15mins to block Cl^−^ uptake by the basolateral Na^+^/K^+^/2Cl^−^ cotransporter Nkcc1 followed by co-treatment with CCH for 15mins to activate Cl^−^ efflux via Ca_2+_-activated Cl^−^ channels (e.g., Ano1). The treatment lowers intracellular Cl^−^ concentration sufficiently to facilitate activity of the basolateral anion exchanger 2 (Ae2). In experiments to acutely block Cftr to increase pH_i_, WT enteroids were treated for 1hr with a combination of 10µM Cftr_inh_-172 and 20µM GlyH101 dissolved in DMSO to evaluate changes in Dvl2-EGFP association with the plasma membrane. Measurements of Dvl2-EGFP intensity were performed by an observer blinded to treatment and genotype.

### Materials

EGF, noggin and Wnt3a were obtained from R&D Systems (Minneapolis, MN). Recombinant Rspondin1 was isolated as described previously^38^.

### Statistics

Cumulative data are reported as the mean ± SE. Intracellular pH data are the average of individual Lgr5-EGFP or crypt-base columnar cells in crypts from WT or Cftr KO enteroids. For juxtamembrane Dvl2-EGFP measurement, individual mouse averages are shown for EGFP intensity which are derived from the average of 1–3 crypt-base columnar cells/crypt from 2–4 crypts of both passage 1 and 2 enteroids. Significant differences between female and male mice were not found for measurements of pH_i_ in Lgr5-EGFP/crypt-base columnar cells or baseline membrane localization of Dvl2-EGF, so these data were combined in the averages. Data between two groups were compared using a two-tailed Student *t*-test or, if not normally distributed with equal variances, by Mann-Whitney Rank Sum test. A probability value of *P*<0.05 was considered statistically significant.

## Results

### Increased intestinal proliferation in Cftr KO crypts is recapitulated in early passage enteroids

Previous in vivo studies of the Cftr KO mouse intestine demonstrated a 34% increase in epithelial proliferation measured by proliferating cell nuclear antigen (PCNA) labeling without a change in apoptosis as compared to WT intestine^14^. To assess intestinal hyperproliferation in our Cftr KO mice in vivo, mitotic cells identified as anti-Phospho Histone H3-positive (PH3+) were enumerated by confocal microscopy using freshly isolated crypts from the proximal small intestine of Cftr KO and WT matched mice (Fig. 1A). Cftr KO crypts had a 52% increase in the number of PH3+ cells per crypt optical cross-section. Mitotic figures were found in both crypt base (< +4 cell position) and the transit-amplifying zone (> +4 position) of each genotype, indicating enhanced proliferation in the ISC and progenitor cell populations. Crypt base columnar cells represent the active ISC population and express the ISC marker Leucine-rich G-protein coupled Receptor 5 (Lgr5)^39^; therefore, Cftr KO heterozygous mice were crossbred with Lgr5-EGFP-ires-CreERT2 (Lgr5-EGFP) mice to compare the average number of Lgr5+ stem cells per crypt between genotypes^30^. Using freshly isolated crypts from WT/Lgr-5-EGFP and Cftr KO/Lgr-5-EGFP matched mice, the number of Lgr5-EGFP-labeled cells was enumerated for each crypt containing at least one Lgr5-EGFP+ cell (Lgr5-EGFP expression is mosaic). As shown in Fig. 1B, the average number of Lgr5-EGFP+ cells per crypt was significantly greater in the Cftr KO/Lgr5-EGFP intestine as compared to WT/Lgr5-EGFP intestine (30.4% increase in mean Lgr5-EGFP+/crypt).

**Fig. 1.**
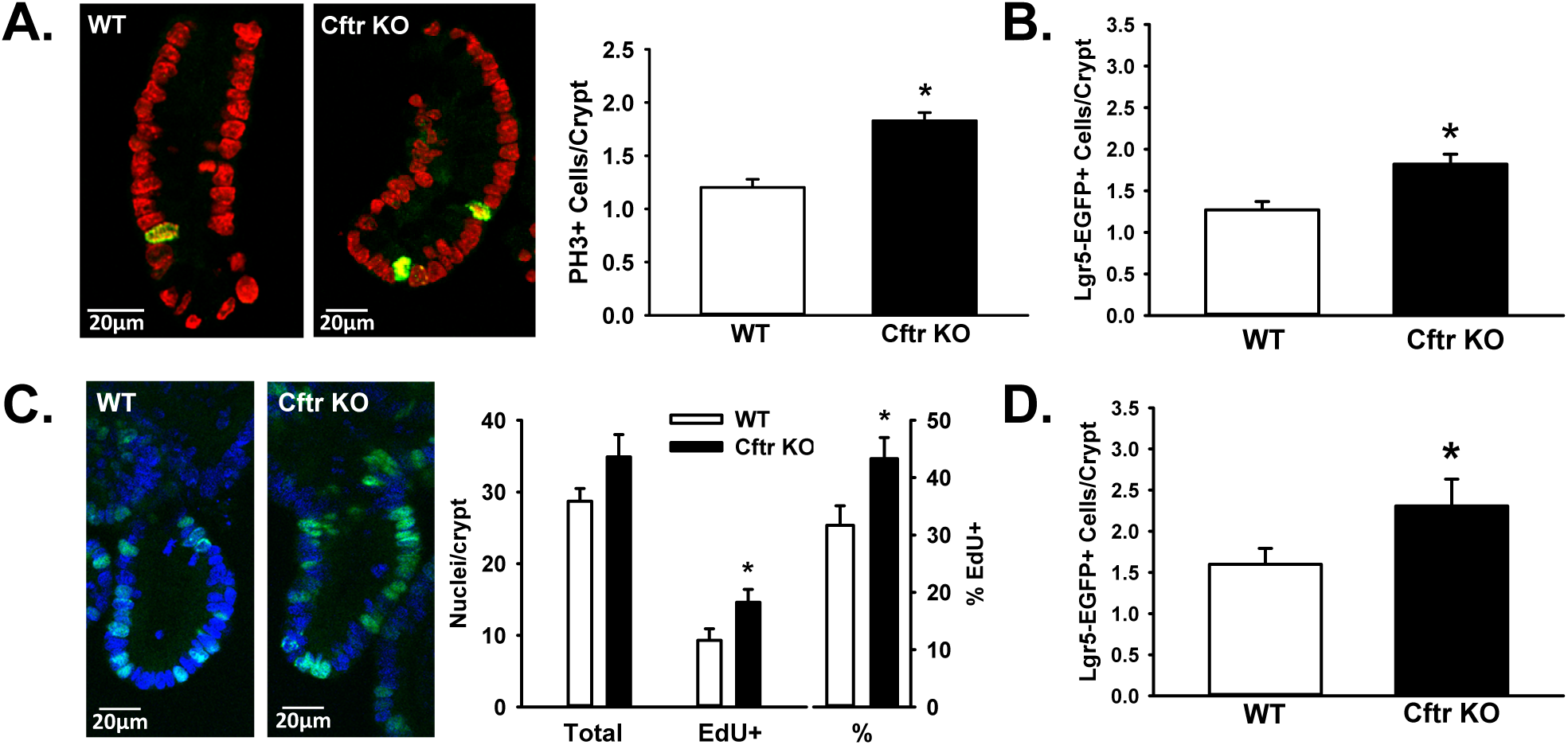
*Increased proliferation in the Cftr KO intestine ex vivo and early passage enteroids. A*: Left, isolated crypts from WT and Cftr KO mice labeled for phospho-Histone H3 (PH3) (green), a marker of mitosis. Nuclei are labeled with TO-PRO 3 nuclear stain (red). Right, cumulative data showing differences in proliferation rates between freshly-isolated WT and Cftr KO crypts as measured by PH3 immunofluorescence. Data are expressed as number of PH3+ cells/crypt optical cross-section; n=4 WT/Cftr KO pairs (32–38 crypts/genotype) \**P*<0.001. *B*: Cumulative data showing the average Lgr5-EGFP+ stem cell numbers/Lgr5-EGFP+ crypt in freshly-isolated crypts from WT/Lgr5-EGFP and Cftr KO/Lgr5-EGFP small intestine. \**P<*0.02; n= 3 WT/Cftr KO pairs (81–113 Lgr5+ crypts/genotype). *C*: Left, enteroid crypts from WT and Cftr KO mice labeled with EdU for S phase nuclei (green). Nuclei are labeled with DAPI nuclear stain (blue). Right, cumulative data showing differences in proliferation rates between WT and Cftr KO enteroid crypts as measured by EdU-positive (EdU+) nuclei and the percentage of EdU+ (%EdU+ nuclei) per crypt optical cross-section. n=5 WT/Cftr KO pairs (52–48 crypts/genotype). \**P*<0.03. *D*. Cumulative data showing the average Lgr5-EGFP+ stem cell numbers/Lgr5-EGFP+ crypt in enteroid crypts from WT and Cftr KO small intestine. \**P<*0.05, n = 5 WT/Cftr KO pairs (156–171 Lgr5+crypts/genotype).

Murine enteroids are well-differentiated, primary cultures of the small intestinal epithelium that can undergo multiple passages^30^. WT enteroid crypts express functional Cftr activity and have proliferation rates that recapitulate in vivo rates when grown as detailed in the Methods^19^, i.e., 25% growth factor concentration relative to the method of Sato et al.^30^. Enteroids from WT and Cftr KO mice enable imaging of live cellular processes in real-time. Furthermore, enteroids avoid the immediate consequences of the abnormal CF intestinal environment^11,3,12,10^, which includes inflammatory mediators and reparative processes that alter Wnt/β-catenin signaling and ISC proliferation^40,41^.

Using passage 1 and 2 cultures, WT and Cftr KO enteroids were exposed to 5-ethynyl-2'-deoxyuridine (EdU) to enumerate cells in S phase. As shown in Fig. 1C, EdU incorporation revealed a 37% increase in proliferation of Cftr KO enteroid crypts compared to WT, as assessed by increases in the total number of EdU+ cells and %EdU+ cells per crypt cross-section in the Cftr KO enteroid crypts. To estimate ISC proliferation, the number of Lgr5-EGFP+ cells in crypts containing at least one Lgr5-EGFP+ cell were enumerated in passage 1 enteroids from WT/Lgr-5-EGFP and Cftr KO/Lgr-5-EGFP matched mice. As shown in Fig. 1D, Lgr5-EGFP+ cells/crypt were significantly greater in the Cftr KO/Lgr5-EGFP as compared to WT/Lgr5-EGFP enteroids (44% increase in mean Lgr5-EGFP/crypt).

We next asked whether later passages (i.e., passage 4) of Cftr KO enteroids would maintain high rates of proliferation, thereby suggestive of an epithelial-autonomous effect of Cftr ablation. At passage 4, WT and Cftr KO enteroids showed similar proliferation rates as measured by EdU labeling. However, the change was largely due to increased proliferation in the WT enteroid crypts which changed from 31.7±3.4 %EdU+ of total crypt nuclei in early passages to 43±4% EdU+ at passage 4 (*P*<0.002, n=5–6). Cftr KO enteroids averaged 42±4% EdU+ of total crypt nuclei in early passage and 43±2% EdU+ at passage 4 (ns, n=5). Corresponding to the increased proliferation in the WT at passage 4, immunoblot analysis of passage 4 WT enteroids revealed decreased protein expression of Cftr (−50.6% relative to passage 1, P<0.02, n = 4–6 WT mice). These findings are consistent with earlier studies of cell lines expressing human CFTR, which showed growth suppression upon functional hCFTR expression and improved proliferation in cells expressing dysfunctional CFTR^42, 43^. We postulate that cells with low levels of CFTR activity have a growth advantage and dominate in subsequent generations, which apparently extends to successive passages of WT enteroid cultures.

### Increased Wnt/β-catenin signaling in Cftr KO intestinal crypts is recapitulated in early passage enteroids

Wnt/β-catenin signaling is required for the maintenance of the ISC population^28^. With evidence of increased proliferation in both freshly-isolated Cftr KO crypts and enteroids, we asked whether Wnt/β-catenin signaling was associated with increased proliferation in the Cftr KO intestine. Freshly-isolated crypts and enteroids from the same WT and Cftr KO matched mice were compared by immunoblot analysis for levels of active β-catenin, defined as β-catenin dephosphorylated at Ser37 and Thr41^44^, and an immediate down-stream Wnt target gene, lymphoid enhancer-binding factor 1 (Lef1)^45^. As a positive control for enhanced Wnt/β-catenin signaling, WT enteroids were also treated with Wnt3a-conditioned medium for 24hrs before collection. The immunoblot shown in Fig. 2A indicates that both freshly-isolated crypts and enteroids from Cftr KO mice have greater levels of active β-catenin and Lef1 as compared to those from matched WT mice. Densitometric analysis showed statically significant increases in active β-catenin and Lef1 in fresh crypts (Fig. 2B) and in the enteroids (Fig. 2C). The above findings indicate that increased Wnt/β-catenin signaling may contribute to increased ISC proliferation in Cftr KO intestine in vivo and in early passage enteroids.

**Fig. 2.**
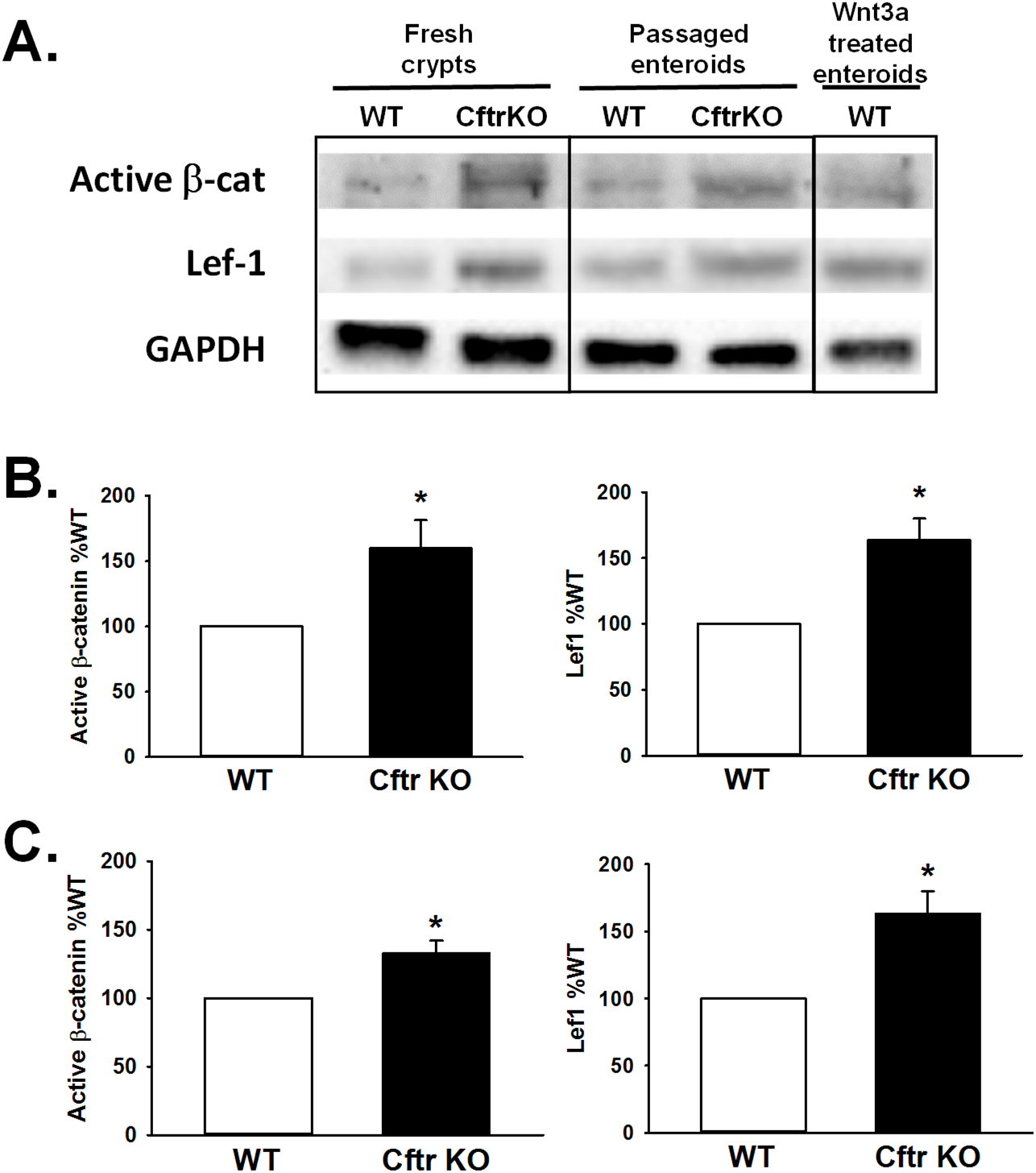
*Increased Wnt/β-catenin signaling in the Cftr KO intestine ex vivo and early passage enteroids. A*: Immunoblot for active β-catenin (dephosphorylated at Ser37/Thr41) and down-stream Wnt target gene Lef1 using freshly isolated crypts (Fresh crypts) and passage 1 enteroids (Passaged enteroids) from a matched pair of WT and Cftr KO mice. Right panel, WT enteroids treated with Wnt3a conditioned medium for 24hrs before collection. *B*. Left, densitometric analysis for immunoblots of active β-catenin from WT and Cftr KO freshly isolated crypts. GAPDH or β-actin used as loading controls. Data represented as % WT; n=11 WT/Cftr KO pairs. \**P*<0.03. Right, densitometric analysis for immunoblots of Lef1 protein expression from WT and Cftr KO isolated crypts. GAPDH or β-actin used as loading control. Data represented as % WT; n=7 WT/Cftr KO pairs. *C*: Left, densitometric analysis for immunoblots of active β-catenin from WT and Cftr KO enteroids. GAPDH or β-actin used as loading controls. Data represented as % WT; n=5 WT/Cftr KO pairs. \**P*<0.02. Right, densitometric analysis for immunoblots of Lef1 protein expression in WT and Cftr KO enteroids. GAPDH or β-actin used as loading controls. Data represented as % WT; n=4 WT/Cftr KO pairs. \**P*<0.03.

### Cftr expression and function in WT Lgr5+ intestinal stem cells

CFTR possesses an intestinal-specific enhancer element in intron 1 that is positively regulated by transcription factor 4 (TCF4) and is therefore a target of Wnt signaling^46^. Based on finding increased Lgr5-EGFP+ stem cell numbers in the Cftr KO intestine (Fig. 1B, 1D) and evidence that CFTR functions as a growth suppressant/tumor suppressor^13, 42, 43^, we asked whether Cftr is expressed and active in ISCs. Different expression levels of SOX9 transcription factor modulate intestinal stem cell proliferation and serve as a marker of crypt cell type^36^. Fluorescence-activated cell sorting of intestinal epithelium from Sox9^EGFP^ transgenic mice have shown that low-expressing Sox9^EGFP^ (Sox9^EGFPLo^) cells are CBCs that are enriched in Lgr5 and form enteroids in culture^36, 47^. Therefore, intestinal epithelial cells from Sox9^EGFP^ mice were sorted into expression fractions and immunoblotted for Cftr. As shown by the immunoblot in Fig. 3A, the Sox9^EGFPLo^ cell fraction had a significant amount of Cftr relative to the Sox9^EGFPSubLo^ fraction, representing transit-amplifying progenitor cells, and the Sox9^EGFPNeg^ fraction, representing the differentiated cell types including enterocytes, goblet cells and Paneth cells.

**Fig. 3.**
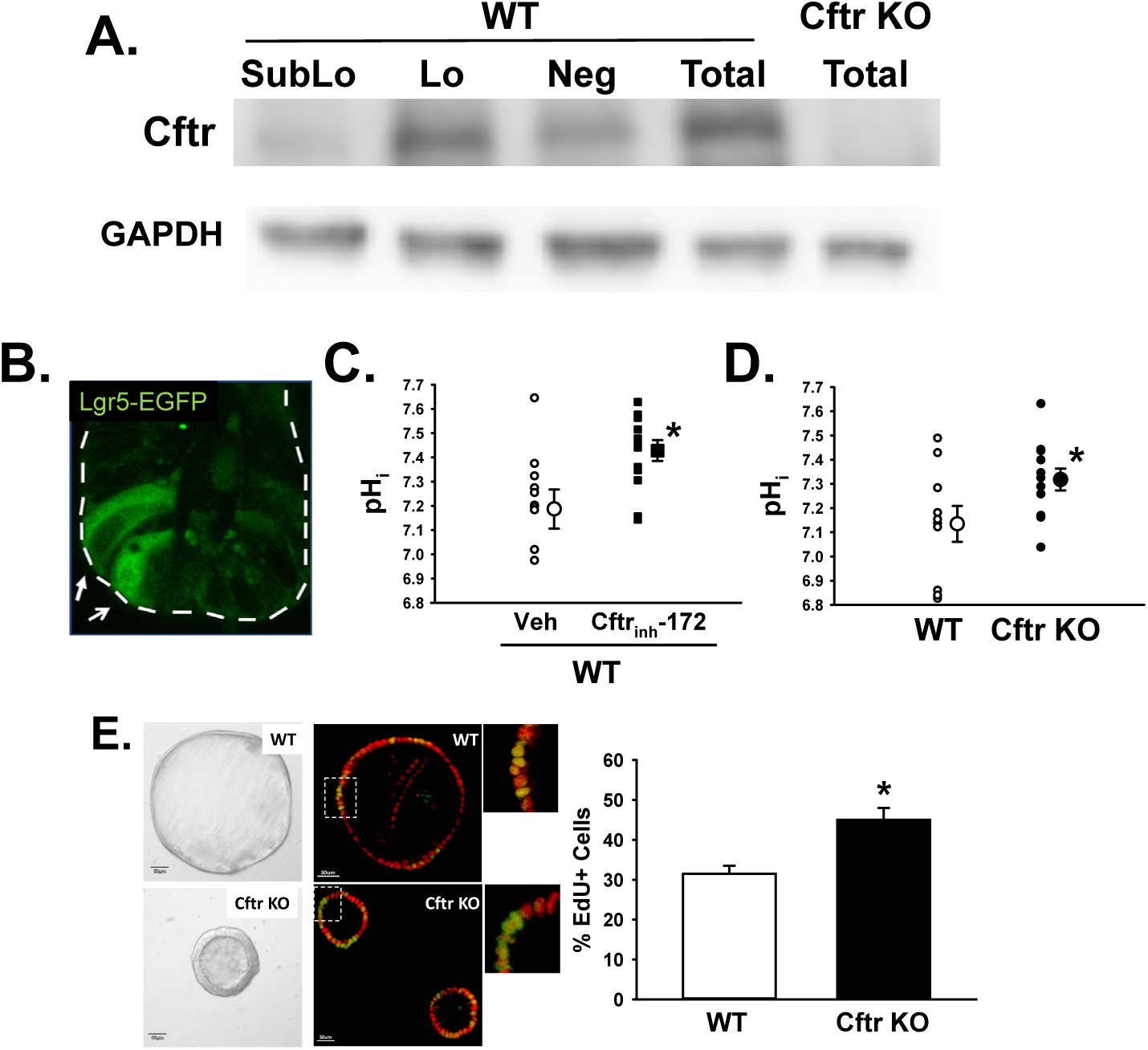
*Expression and function of Cftr in small intestinal stem cells. A*. Immunoblot for Cftr in flow cytometry cell fractions of small intestinal epithelium isolated from Sox9^EGFP^ mice. Sublo, sub-low Sox9^EGFP^-expressing cell fraction (transit-amplifying progenitors); Lo, low Sox9^EGFP^-expressing cell fraction (crypt-base stem cells, Lgr5+); Neg, negative for Sox9^EGFP^ expression cell fraction (enterocyte, goblet, Paneth); Total, total small intestinal epithelium. Cftr KO total epithelium, negative control. GAPDH, loading control. Representative of 3 Sox9^EGFP^ mice. *B*. Confocal micrograph of Lgr5-EGFP-expressing intestinal stem cells in crypt epithelium (white arrows). White dashed line outlines crypt. *C*. Intracellular pH of Lgr5-EGFP+ cells in WT/Lgr-5-EGFP enteroids treated with either DMSO (Veh, small circles) or Cftr_inh_-172 (10µM for 1hr, small squares). Treatment average, large circle or square. \**P*<0.02 vs. Veh, n = 10–13 Lgr5-EGFP cells from 7–8 enteroids cultured from 6 WT/Lgr5-EGFP mice. *D*. Intracellular pH of Lgr5-EGFP+ cells in enteroids from matched WT/Lgr-5-EGFP (open small circles) and Cftr KO/Lgr-5-EGFP (filled small circles) mice. Average shown as large circles. \**P*<0.04, n = 10–12 Lgr5-EGFP cells from 6 enteroids from 3 WT/Lgr-5-EGFP and Cftr KO/Lgr-5-EGFP matched mice. *E*. Left, ISC-enriched enterospheres from WT and Cftr KO matched mice and a photomicrograph showing EdU positive nuclei (green) and nuclei (red) in an optical cross-section of WT and Cftr KO enterospheres. Right, percentage of EdU-positive nuclei relative to total nuclei in optical cross-sections of WT and Cftr KO enterospheres. (Total number of nuclei/enterosphere were WT = 36.1±3.8; Cftr KO = 46.7±2.0, n = 32 enterospheres/genotype; *P*<0.009). \**P*<0.05 vs. WT. n= 4 WT and Cftr KO matched mice (8 enterospheres/mouse).

To investigate Cftr functional activity in ISCs, we examined the effect of Cftr inhibition or ablation on the intracellular pH (pH_i_) of Lgr5-EGFP+ stem cells using enteroids from matched WT/Lgr5-EGFP and Cftr KO/Lgr5-EGFP mice. Previous studies have shown that acute inhibition of Cftr using the channel blocker Cftr_inh_-172 alkalizes pH_i_ in the crypt epithelium of WT enteroids19. More importantly, pH_i_ of crypt epithelium in Cftr KO enteroids is uncompensated and maintains an alkaline pH_i_19, 20, similar to reports of duodenal villous epithelium from Cftr KO mice^37, 48^ and CFPAC cells, a pancreatic cell line derived from a CF patient^49^. To measure intracellular pH in Lgr5-EGFP+ stem cells (Fig. 3B), enteroids were loaded with the ratiometric, pH-sensitive dye SNARF-5F for confocal microfluorimetry. As shown in Fig. 3C, the pH_i_ in WT Lgr5-EGFP stem cells significantly increased by ~0.2 pH units following a 1 hr treatment with Cftr_inh_-172 (10µM). Moreover, as shown in Fig. 3D, Cftr KO Lgr5-EGFP+ stem cells exhibit an alkaline pH_i_ that is also ~0.2 pH units greater than in WT Lgr5-EGFP+ stem cells. These findings indicate that both acute and chronic loss of Cftr function alkalizes Lgr5-EGFP+ intestinal stem cells, indicative of a pH_i_ regulatory role by Cftr anion channel activity in WT ISCs.

Primary culture of dissociated small intestinal crypt epithelium with exogenous Wnt3a (100 ng/mL) results in the formation of ISC–enriched epithelial spheroids (enterospheres)^50–54^. In examining Cftr stem cell activity, we asked whether ISC-enriched enterospheres from WT and Cftr KO would exhibit differences in proliferation consistent with our study enumerating Lgr5-EGFP+ ISCs/crypt (Figs 1B, 1D). As shown in Fig. 3E (left), WT enterospheres formed enlarging monolayer cysts, suggesting Cftr-mediated fluid secretion. In contrast, Cftr KO enterospheres had much smaller luminal cavities and slow cyst expansion with time in culture, as noted previously in studies of human CF colon organoids^55^. EdU and confocal optical cross-section studies were used to examine proliferation rates in WT and Cftr KO enterospheres. As shown in Fig. 3E (right), Cftr KO enterospheres had a 38% greater rate of proliferation relative to WT enterospheres. These findings demonstrate a role of Cftr as growth suppressant in an enriched ISC population.

### Intracellular alkalinity and membrane association of Dvl2-EGFP in Cftr KO crypt-base columnar cells

Different facets of Wnt signaling play major roles in cellular processes of proliferation, cell migration and polarity^27^. The signaling mediator Disheveled (Dvl) is critical for downstream activation of both canonical and non-canonical Wnt signaling pathways through its recruitment to Frizzled (Fz) membrane receptors^56^. The formation of a stable Dvl-Fz association involves both the binding of the PDZ domain of Dvl to the PDZ binding domain of Fz and the binding of the polybasic DEP domain of Dvl to negatively charged phosphatidic acid (PA) and phosphatidylglycerol (PG) at the inner leaflet of the plasma membrane26,57. The latter process is both charge- and pH_i_- dependent, wherein a more alkaline pH_i_ facilitates electrostatic attraction of the polybasic DEP domain to the cell membrane. Importantly, the three major domains of Dvl are conserved across species, including the PDZ and DEP domains^27^. Based on evidence of alkalinity and increased proliferation in Cftr KO ISCs, we asked whether Dvl in Cftr KO CBCs associates more closely with the plasma membrane as a means to facilitate Wnt/β-catenin signaling.

Subcellular localization of the major Dvl isoform in mice, Dvl2, was imaged by confocal microscopy in live WT and Cftr KO enteroids from transgenic mice that express Dvl2-EGFP (native Dvl2 KO) and the Rosa^mT/mG^ plasma membrane reporter (Fig.4A). Measurements of Dvl2-EGFP proximity to the cell membrane were limited to the supranuclear region, i.e., between the apical pole of the nucleus and the apical membrane, due to the close apposition of the nucleus/nuclear membrane to the plasma membrane in the basal portion of the cell. To evaluate whether pH_i_ differences between WT/Dvl2-EGFP/Rosa^mT/mG^ and Cftr KO/Dvl2-EGFP/Rosa^mT/mG^ CBC cells were present in this cell region, high resolution SNARF-5F confocal microfluorimetry measured pH_i_ in the supranuclear region of non-granulated CBC cells, serving as proxy for Lgr5+ ISCs^30^. As shown in Fig. 4B, Cftr KO enteroids exhibited a significantly more alkaline pH_i_ in the supranuclear region of CBCs as compared to WT, therefore confirming pH_i_ differences in this region. As depicted in the model shown in Fig. 4C, the membrane proximity of Dvl2-EGFP to the supranuclear lateral membrane of a CBC was measured as the EGFP intensity within a two-pixel distance (1.15µm) interior to the lateral and medial mTomato plasma membranes (relative to crypt axis) from the apical pole of the nucleus to the apical membrane. As shown in Fig. 4D, left, the Cftr KO CBCs exhibited greater (~two-fold) juxtamembrane Dvl2-EGFP intensity at the supranuclear lateral plasma membranes than was found in WT CBCs. Interestingly, as shown in Fig. 4D, right, the WT CBCs showed a polarized distribution of Dvl2-EGFP with significantly greater juxtamembrane Dvl2-EGFP intensity at the lateral-Lateral membrane than at the medial-Lateral cell membrane. The juxtamembrane Dvl2-EGFP intensity at medial-Lateral membrane was not different from the average EGFP intensity of all supranuclear pixels, i.e., no detectable association. In contrast, the juxtamembrane Dvl2-EGFP intensity at the lateral-Lateral membrane in the Cftr KO CBCs was similar to that of WT but significantly greater than WT at the medial-Lateral membrane, indicating a less polarized distribution of Dvl2 at the lateral plasma membrane in Cftr KO CBCs.

**Fig. 4.**
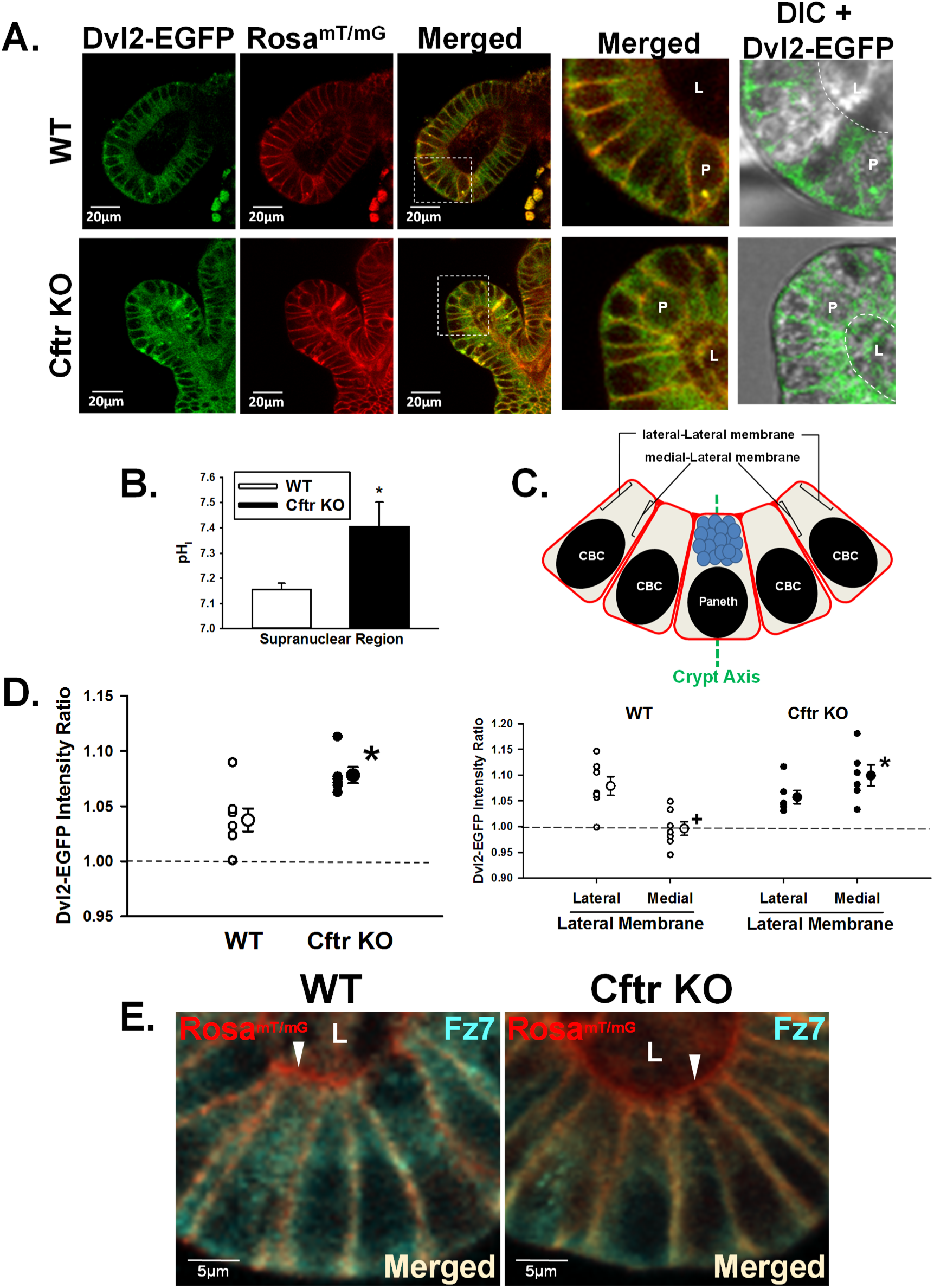
*Increased Dvl2-EGFP localization at the plasma membrane in Cftr KO crypt-base columnar cells. A.* Representative images of WT/Dvl2-EGFP/Rosa^mT/mG^ (top) and Cftr KO/Dvl2-EGFP/Rosa^mT/mG^ (*bottom*) enteroid crypts showing Dvl2-EGFP (green), Rosa^mT/mG^ (red), and Merged images. Magnified Merged and differential interference contrast merged with Dvl2-EGFP (DIC+Dvl2-EGFP) images of crypt base (from white boxes), were magnified (3.6X). P, Paneth cell; L, crypt lumen. White dashed lines outline apical membrane orient towards the crypt lumen. *B.* Cumulative pH_i_ data for the supranuclear region of crypt-base columnar cells (CBCs) in WT and Cftr KO enteroids. \**P*<0.03 vs. WT; n = 4–8 crypts from 4 WT and Cftr KO matched mice. *C*. Crypt base model depicting the measurement by confocal microfluorimetry of Dvl2-EGFP intensity two pixels interior to the lateral-Lateral Membrane (lateral to central crypt axis) and medial-Lateral Membrane (medial to central crypt axis) plasma membrane of CBCs. Green dashed line, central crypt axis. Brackets indicate sites of measurement on the supranuclear lateral plasma membranes. CBC, crypt-base columnar cell. Paneth, Paneth cell. *D. Left*, cumulative data showing the average Dvl2-EGFP intensity ratio within 1.15µm of the CBC supranuclear plasma membrane of individual mice (small circles) and overall average (large circles) for WT/Dvl2-EGFP/Rosa^mT/mG^ (WT) and Cftr KO/Dvl2-EGFP/Rosa^mT/mG^ (Cftr KO) sex-matched mouse pairs (n= 7 and 6 mice, respectively). Dashed line, indicates the average Dvl2-EGFP pixel intensity for the entire supranuclear region of CBC which has been set to 1.0 (Ave. intensity = 59.5±8.8 for WT and 88.5±15.9 for Cftr KO, ns, n = 7–6 mice, respectively (average of 1–3 CBCs/crypt from 2–4 crypts of passage 1 and 2 enteroids). \**P*<0.01 vs WT plasma membrane. *Right,* Cumulative data showing the average Dvl2-EGFP intensity ratio at the individual supranuclear lateral-Lateral and medial-Lateral plasma membranes of individual mice (small circles) and overall average (large circles) for WT/Dvl2-EGFP/Rosa^mT/mG^ (WT) and Cftr KO/Dvl2-EGFP/Rosa^mT/mG^ (Cftr KO) sex-matched mouse pairs (n= 7 and 6 mice, respectively). Dashed line, indicates the average Dvl2-EGFP pixel intensity for the entire supranuclear region of CBC which has been set to 1.0. +*P*<0.002 vs WT lateral-Lateral cell membrane. \**P*<0.001 vs. WT medial-Lateral cell membrane. *E*. Immunofluorescence image of Frizzled 7 (Fz7, cyan), plasma membrane (red) and merged (light yellow) of CBCs in WT and Cftr KO enteroid crypts. Representative of 3 WT-Cftr KO mouse pairs. L, lumen. White arrowheads denote lack of Fz7 staining at apical membrane (red).

The Dvl2-EGFP intensity ratio shown in Fig. 4D are averages of WT and Cftr KO matched mice. In the analysis, we found Dvl2-EGFP intensity ratio between individual enteroid crypts (consisting of the average of 1–3 CBCs/crypt) exhibited greater variability than between individual mice of the same genotype. However, the average Dvl2-EGFP intensity ratio at the lateral cell membranes of WT and Cftr KO crypts was nearly identical to the per mouse averages shown in Fig. 4D, right (WT = 1.04±0.01 vs. Cftr KO = 1.08±0.01 Intensity Ratio, n = 18 and 13 crypts, respectively, *P*<0.02). Thus, enteroid crypt averages from paired WT-Cftr KO mice also provide a robust comparison of the juxtamembrane Dvl2-EGFP intensity ratio between the two genotypes.

To determine whether increased localization of Dvl2 at the supranuclear lateral plasma membrane was positioned to facilitate Wnt/β-catenin signaling, immunofluorescence studies evaluated the plasma membrane localization of the primary Wnt receptor for canonical Wnt/β-catenin signaling in ISC, i.e., Frizzled 7 (Fz7)^58^. As shown in Fig. 4E, the distribution of Fz7 appeared uniform along the entire lateral plasma membrane of CBC cells in both WT and Cftr KO enteroids. Fz7 was not localized to the apical (lumen-facing) membrane of CBCs. Thus, the supranuclear region of live CBCs is appropriate for evaluating Dvl2-EGFP membrane association and its potential for facilitating Wnt/β-catenin signaling in WT and Cftr KO enteroids.

### Neutralization of plasma membrane charge in Cftr KO enteroids reduces Dvl2 proximity to the plasma membrane

The DEP domain of Dvl binds to acidic lipid headgroups of phosphatidic acid and phosphotidlylglycerol at physiological pH^26^. In HEK293T cells, neutralization of negatively-charged phospholipids with the cationic lipid sphingosine reduced Dvl-Fz association by interfering with the electrostatic interaction of the Dvl DEP domain with the inner leaflet of the plasma membrane^26^. To examine the effect of neutralizing the negatively-charged inner leaflet phospholipids in Cftr KO CBCs, Dvl2-EGFP intensity at the supranuclear plasma membrane was measured in Cftr KO enteroids treated for 1hr with 75µM sphingosine or vehicle (Fig. 5A, left). As shown in Fig. 5A, right, Dvl2-EGFP intensity at the CBC supranuclear plasma membrane was significantly reduced in sphingosine-treated enteroids as compared to vehicle control.

**Fig. 5.**
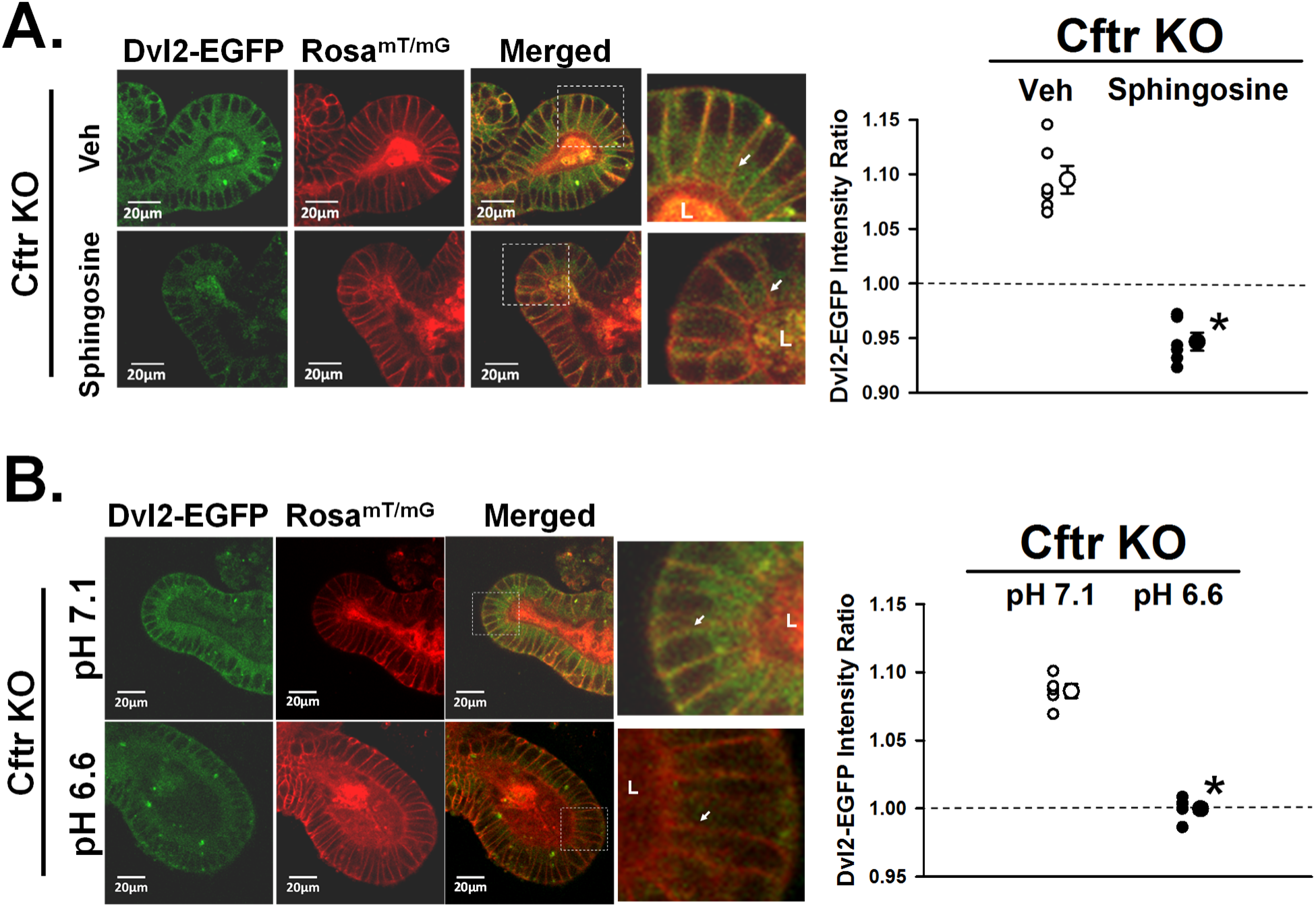
*Dvl2-EGFP proximity to the plasma membrane is charge- and pH^i^–dependent in Cftr KO crypt-base columnar cells*. A. Left, representative images of ETOH vehicle-(Veh, top) and sphingosine-treated (Sphingosine, bottom) Cftr KO/Dvl2-EGFP/Rosa^mT/mG^ enteroid crypts showing Dvl2-EGFP (green), Rosa^mT/mG^ (red) and merged images. Magnified merged image (2.5 X) of crypt base (from white boxes). L, crypt lumen; lateral cell membrane for comparison (white arrows). Right, cumulative data showing the average Dvl2-EGFP intensity ratio within 1.15µm of the CBC supranuclear plasma membrane of individual mice (small circles) and overall average (large circles) for Veh- and Sphingosine-treated Cftr KO/Dvl2-EGFP/Rosa^mT/mG^ (Cftr KO) mice (n=6). Dashed line, indicates the average Dvl2-EGFP pixel intensity for the entire supranuclear region of CBC which has been set to 1.0. \**P*<0.001 vs. Veh. Averaged data for each mouse represents measurements from 1–3 CBCs/crypt from 2–4 crypts from passage 1 and 2 enteroids. *B*. Left, representative images of Cftr KO enteroids cultured for 48hr in Medium pH 7.1 (top) and Medium pH 6.6 (bottom) showing Dvl2-EGFP (green), Rosa^mT/mG^ (red) and merged images. Magnified merged image (4.1X) of crypt base (from white boxes). L, crypt lumen; lateral cell membrane for comparison (white arrows). Right, cumulative data showing the average Dvl2-EGFP intensity ratio within 1.15µm of the CBC supranuclear plasma membrane of individual mice (small circles) and overall average (large circles) for pH 7.1- and pH 6.6-treated enteroids from Cftr KO/Dvl2-EGFP/Rosa^mT/mG^ (Cftr KO) mice (n=4). Dashed line, indicates the average Dvl2-EGFP pixel intensity for the entire supranuclear region of CBC which has been set to 1.0. \**P*<0.001 vs. pH 7.1 medium. Averaged data for each mouse represents measurements from 1–3 CBCs/crypt from 2–4 crypt from passage 1 and 2 enteroids.

Sphingosine significantly reduced Dvl2-EGFP membrane association at both the lateral-Lateral and medial-Lateral cell membranes of the Cftr KO CBCs (Veh lateral-Lateral = 1.09±0.02 and medial-Lateral = 1.10± 0.01 vs. Sphingosine lateral-Lateral = 0.92±0.02 vs. medial-Lateral = 0.97±0.01 Intensity Ratio; vs. Veh lateral-Lateral membrane *P*< 0.001; vs. Veh medial-Lateral membrane *P*<0.001). A comparison of enteroid crypt averages for vehicle and sphingosine treatments yielded similar results (Veh = 1.10±0.01 vs. Sphingosine = 0.95±0.01 Intensity Ratio, n = 11 crypts each, *P*<0.000000001). Prolonged sphingosine treatment (2–3hrs) proved toxic in that enteroids began to dissociate with cell rounding, therefore, we did not attempt to measure active β-catenin protein under this condition. Nonetheless, the above findings indicate that Dvl2 association with the plasma membrane is charge-dependent in Cftr KO CBC cells.

### Reduction of pH_i_ in Cftr KO enteroid CBCs reduces Dvl2 plasma membrane association

Alkalinity reduces proton interaction with the negatively-charged domains of membrane phospholipids allowing for greater interaction with Dvl2 DEP domain^26, 59^. Therefore, the pH_i_ in Cftr KO enteroid crypts was manipulated by culturing Cftr KO/Dvl2-EGFP/Rosa^mT/mG^ enteroids for 3 days in pH 7.1 medium followed by 2 days in medium at either pH 6.6 or maintained in pH 7.1 medium (controls). In the pH 6.6 medium, the pH_i_ was reduced by −0.31 ± 0.06 pH units relative to pH 7.1 medium (*P*<0.0002, n= 13 CBC from 2–4 enteroids from 3 matched WT and Cftr KO mice). Overt cell/enteroid morphology were unaffected by pH 6.6 medium (Fig. 5B, left). Cftr KO enteroids cultured in pH 6.6 medium, as compared to pH 7.1 medium, exhibited reduced Dvl2-EGFP intensity at the supranuclear lateral plasma membranes of CBCs (Fig. 5B, right). Both the lateral-Lateral and medial-Lateral membranes showed reduced Dvl2-EGFP intensity in the acidified Cftr KO CBCs (pH 7.1: lateral-Lateral = 1.08±0.01 and medial-Lateral = 1.10± 0.01; pH 6.6: lateral-Lateral = 1.00±0.03 vs. medial-Lateral = 1.00±0.03 Intensity Ratio *P*< 0.03 and *P*<0.01, respectively). Similar results were yielded by comparison of crypt averages for the two conditions (pH 7.1 = 1.09±0.01 vs. pH 6.6 = 0.99±0.01 Intensity Ratio, n = 11 and 7 crypts, respectively, *P*<0.00000002). Measurements of Cftr KO enteroids exposed to the pH 6.6 medium showed a significantly reduced active β-catenin as compared to pH 7.1 medium (as % of pH 7.1 (100%): pH 6.6 = 64.9±5.1%, P<0.03, enteroids from n=5 Cftr KO mice). Acidic conditions influence cell proliferation in several ways^60^; therefore, proliferation of Cftr KO enteroids maintained in pH 6.6 medium was not investigated.

### Acute pharmacological manipulation of Dvl2-EGFP plasma membrane association in CBCs

Recent studies have shown that the alkaline pH_i_ of crypt epithelium in Cftr KO enteroids can be acutely reduced by pharmacological reduction of intracellular [Cl^−^] ([Cl^−^]_i_), which facilitates basolateral anion exchanger 2 normalization of pH_i_ 20. To further assess the relationship between pH_i_ and Dvl2-EGFP membrane association, Cftr KO/Dvl2-EGFP/Rosa^mT/mG^ enteroids were treated for 30min with a sequential combination of 50µM bumetanide (15min) to inhibit Cl^−^ uptake by the Na^+^/K^+^/2Cl^−^ cotransporter NKCC1, followed by 100µM carbachol (15min) to induce epithelial Cl^−^ secretion by Ca^2+^-activated Cl^−^ channels (e.g., Ano1). As shown in Fig. 6A, pharmacological treatment to normalize pH^i^ by reducing [Cl^−^]^i^ significantly reduced Dvl2-EGFP association at the supranuclear lateral plasma membranes in Cftr KO CBCs. Both the lateral-Lateral and medial-Lateral membranes had significantly reduced Dvl2-EGFP membrane association in the treated Cftr KO CBCs (Veh: lateral-Lateral = 1.08±0.02 and medial-Lateral = 1.10± 0.02; Bumet-CCH: lateral-Lateral = 0.97±0.04 vs. medial-Lateral = 1.02±0.01 Intensity Ratio, *P*< 0.03 and *P*<0.03, respectively). In a converse experiment, WT enteroids were treated with a combination of 10µM Cftr_inh_-172 and 20µM GlyH-101 for 1hr to inhibit Cftr Cl^−^ and HCO^3^- conductance at both extra- and intracellular sites, respectively^61^. As shown in Fig. 6B, pharmacological inhibition of Cftr to increase WT ISC pH_i_ significantly increased Dvl2-EGFP association with the supranuclear lateral cell plasma membrane to a level similar to Cftr KO CBCs (see Fig. 4D). The increase in Dvl2-EGFP membrane association primarily occurred at the medial-Lateral plasma membrane in the WT enteroids treated with Cftr inhibitors (Veh: lateral-Lateral = 1.05±0.03 and medial-Lateral = 1.02±0.01; Cftr inhibitors lateral-Lateral = 1.09±0.01 and medial-Lateral = 1.10±0.01 Intensity Ratio; ns and *P*<0.0005, respectively). Both of the above studies were paralleled by crypt averages for the two experiments (Veh = 1.09±0.01 vs. Bumet-CCH = 0.99±0.02 Intensity Ratio, n = 6 and 8 crypts, respectively, *P*<0.003; Veh = 1.03±0.01 vs. Cftr_inh_172-GlyH 101 = 1.09± 0.01 Intensity Ratio, n = 6 crypts each, *P*<0.009). Since both manipulations of pH_i_ were acute, it was not expected that downstream accumulation of active β-catenin would be sufficient to warrant measurement.

**Fig. 6.**
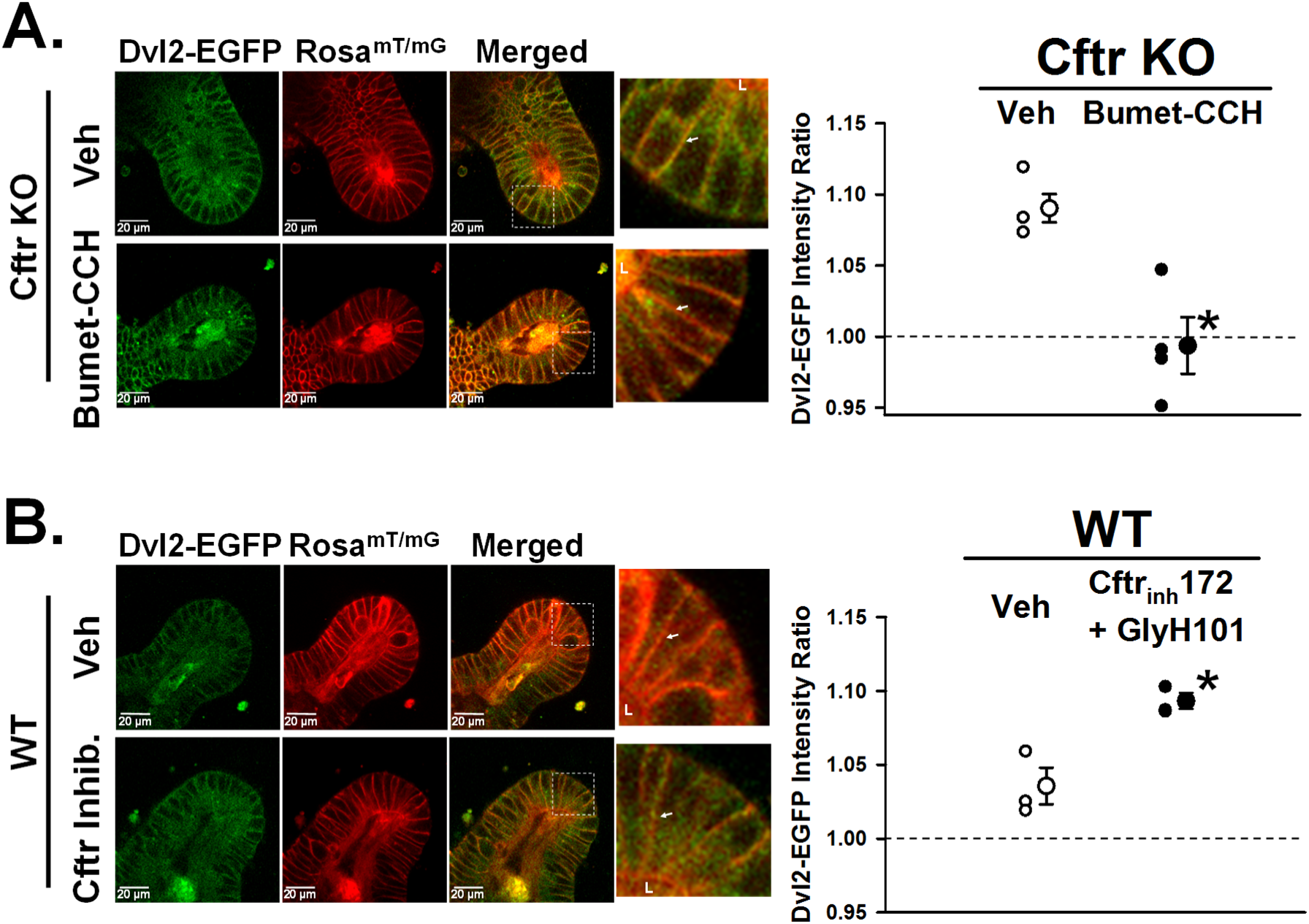
*Pharmacological manipulation of Dvl2-EGFP plasma membrane association. A*. Left, representative images of DMSO vehicle- (Veh, top) and bumetanide+carbachol-treated (Bumet-CCH, bottom) Cftr KO/Dvl2-EGFP/Rosa^mT/mG^ enteroid crypts showing Dvl2-EGFP (green), Rosa^mT/mG^ (red) and merged images. Magnified merged image (3.6 X) of crypt base (from white boxes). L, crypt lumen; lateral cell membrane for comparison (white arrows). Right, cumulative data showing the average Dvl2-EGFP intensity ratio within 1.15µm of the CBC supranuclear lateral plasma membrane of individual mice (small circles) and overall average (large circles). Cftr KO/Dvl2-EGFP/Rosa^mT/mG^ (Cftr KO) mice were treated with Vehicle (Veh, DMSO) or sequential exposure to 50µM bumetanide (Bumet, 15min) followed by 100µM carbachol (CCH, 15min), n= 4 mice. Averaged data for each mouse represents measurements from 1–3 CBCs/crypt from 2–4 crypts from passage 1 and 2 enteroids. Dashed line, indicates the average Dvl2-EGFP pixel intensity for the entire supranuclear region of CBC which has been set to 1.0. \**P*<0.005 vs. Vehicle. *B*. Left, representative images of DMSO vehicle- (Veh, top) and Cftrinh-172+GlyH-101-treated (Cftr Inhib., bottom) WT/Dvl2-EGFP/Rosa^mT/mG^ enteroid crypts showing Dvl2-EGFP (green), Rosa^mT/mG^ (red) and merged images. Magnified merged image (3.8 X) of crypt base (from white boxes). L, crypt lumen; lateral cell membrane for comparison (white arrows). Right, cumulative data showing the average Dvl2-EGFP intensity ratio within 1.15µm of the CBC supranuclear lateral plasma membrane of individual mice (small circles) and overall average (large circles). WT/Dvl2-EGFP/Rosa^mT/mG^ (WT) mice were treated with Vehicle (Veh, DMSO) or 10µM Cftrinh-172 plus 20µM GlyH-101 for 1hr, n= 3 mice. Averaged data for each mouse represents measurements from 1–3 CBCs/crypt from 2–4 crypts from passage 1 and 2 enteroids. Dashed line, indicates the average Dvl2-EGFP pixel intensity for the entire supranuclear region of CBC which has been set to 1.0. \**P*<0.013 vs. Vehicle.

## Discussion

Increased proliferation of the stem cell compartment is a potential risk factor for intestinal cancer^15^. Approximately three mutations occur with every division of a normal stem cell, therefore augmented stem cell proliferation raises the potential for a DNA replication error leading to ‘driver’ cancer mutations^16^. Cancer genome sequencing and epidemiological studies suggest that stochastic DNA replication errors are responsible for two-thirds of the mutations in human cancers^15, 16^. One of the less commonly recognized manifestations of cystic fibrosis intestinal disease is an increased risk for gastrointestinal cancer. Surprising for a relatively young population, previous studies have shown that CF patients have a 6–10 fold increased risk for GI cancer, whereas the risk of non-digestive tract cancers in CF patients is similar to the general population^9, 62^. The risk of gastrointestinal cancer increases to 20-fold in patients 20–29 years of age and includes a significantly greater number of small intestinal tumors than in the non-CF population^9^. In accordance with increased GI cancer risk in CF patients, a recent study provided evidence that CFTR is a tumor suppressor gene in human intestinal cancer^13^. The same study also demonstrated a strong propensity of CF mice for spontaneous development of intestinal tumors when aged to 1 year with a penetrance of 60% as compared to 0% in WT mice. The majority of tumors in CF mice were in the small intestine. One factor to consider here is that stem cell proliferation in the human colon exceeds that in the small bowel, whereas the inverse apparently holds true for mice^15^.

Another factor deserving consideration is the propensity for intestinal cancer versus other cell/organ types in CF, which is not the case with many tumor suppressor mutations or knockout models^63^. The diurnal and postprandial activity of Cftr in the alimentary tract is dynamic which likely accentuates its role in pH_i_ regulation and thus Wnt/β-catenin signaling in these epithelia. Unlike familial adenomatous polyposis (FAP) or Lynch syndrome that carry greater intestinal cancer risk due to mutations directly involved in β-catenin signaling or DNA mismatch repair^64, 65^, Cftr’s role in growth suppression as outlined in this study is indirect and subject to known compensatory forces in the regulation of pH_i_20. Therefore, the propensity for gastrointestinal cancer in CF likely involves the combination of a life-long increase in ISC proliferation and the CF intestinal environment which presents additional cancer risk factors including low-grade inflammation^4, 5, 11^, small bowel bacterial overgrowth^3, 12^, dysbiosis^66–68^ and goblet cell metaplasia^69^. With advances in CF respiratory therapy leading to increased life expectancy and the current prevalence of lung transplantation in CF patients^8^, there is a need to better understand the causes of intestinal hyperproliferation in the adult CF intestine and its relationship with cancer incidence.

Increased ISC proliferation in the Cftr KO intestine was maintained in early passage primary enteroid culture, suggestive of an inherent epithelial phenotype. The Cftr KO proliferative phenotype was apparent through culture passages 1 and 2, but proliferation rates of WT and Cftr KO enteroid crypts converged by passage 4. However, matched WT enteroids at passage 4 assumed a CF-like phenotype with decreased Cftr expression and increased proliferation, ostensibly due to selection against slower growth by cells (ISCs) expressing higher amounts of Cftr^42, 43^. In vivo, the CF intestinal environment may promote epithelial proliferation through disease factors known to increase the risk for GI cancer. One factor is intestinal inflammation; however, studies of small intestinal inflammation in both CF patients and CF mice indicate a relatively mild presentation. Neutrophil and mononuclear cell infiltration of the submucosa are moderate without overt mucosal damage^5, 11^. Gene expression studies indicate immune activation in the small intestine of CF patients^4, 5^. In CF mice, changes in innate immunity genes were also noted, but minimized by therapeutic intervention to prevent bowel obstruction by continuously maintaining the mice on a polyethylene glycol (PEG) osmotic laxative in the drinking water^70^. CF mice in the present investigation and in the study by Than et al. (showing a high incidence of GI tumors) were both maintained on the PEG laxative, which may reduce the contribution of inflammation-induced genetic/epigenetic changes. Together with evidence that Cftr is expressed and functional in murine Lgr5+ ISCs, we tentatively conclude that hyperproliferation in the CF intestine is a consequence of losing the growth suppressive function of Cftr. However, without further investigation, it is difficult to eliminate the possibility that growth stimulation in the Cftr KO enteroids includes epigenetic changes secondary to the CF intestinal environment that are carried into culture.

Wnt/β-catenin signaling is required for proliferation and maintenance of small intestinal stem cells^28^. We found that Wnt/β-catenin signaling, as indicated by active β-catenin and down-stream Lef1 protein amounts, is increased in both freshly isolated crypts and early passage enteroids of Cftr KO mice. This finding is consistent with increased numbers of Lgr5+ stem cells in the intestine and enteroids of Cftr KO/Lgr5-EGFP mice. Further studies revealed Cftr protein expression in sorted SOX9^EGFPLo^ cells, an alkaline pH_i_ in Lgr5+-EGFP ISCs in the absence of Cftr activity and greater proliferation in ISC-enriched enterospheres from Cftr KO mice as compared to WT. This evidence supports previous studies identifying CFTR as a down-stream target for the terminal effector of Wnt/β-catenin signaling, Tcf4, and suggests that Cftr may play a role in maintaining the balance between intestinal proliferation and differentiation^46^. As a direct target gene, Cftr may also modulate ISC proliferation through actions on the cell cycle. In T and B lymphocyte cell lines, it has been shown that a CFTR-dependent Cl^−^ permeability is increased during the G1 phase of the cell cycle which could be augmented by increasing intracellular cAMP^71^. This coordinates nicely with evidence in other cell types that increased cAMP and reduced intracellular [Cl^−^] during G1 leads to cell cycle arrest by a p21-dependent mechanism^72, 73^. However, potential involvement of Cftr as a growth suppressant in ISC by cell cycle regulation does not elucidate the pathway of increased Wnt/β-catenin signaling in the crypts of the Cftr KO intestine.

The pathogenesis connecting dysregulated Wnt/β-catenin signaling to the absence of an apical membrane anion channel is unlikely to be direct. Although previous studies have shown CFTR to interact with several proteins thereby affecting various cell processes74, 75, the undisputed function of CFTR is to provide epithelial Cl^−^ and HCO_3_- permeability. Loss of this permeability in CF disease has been associated with dehydration and reduced pH of the airway and intestinal surfaces, increased mucus viscosity, and deficits of innate immunity^76–81^. Another cellular consequence of CFTR loss is dysregulation of pH_i_ in the alkaline range, which was recently shown to be dependent on the combined intracellular retention of Cl^−^ and HCO_3_- 20. Alkaline pH_i_ is conducive to cell proliferation by positive effects on cell cycle transitions and DNA replication^21, 22^ and is known to facilitate Wnt signaling by increasing the interaction of Dvl with the inner plasma membrane to stabilize binding with the Wnt receptor, Fz^26^. Dvl possesses a polycationic motif that targets the protein to the negatively charged inner leaflet, similar to the process used for subcellular localization of a variety of polycationic proteins including K-Ras and proteins associated with endosomes (which demonstrate defective recycling in CF cells^82^). Discovered through a genome-wide RNAi screen of *Drosophila* cells, disheveled (Dsh) membrane localization required Na^+^/H^+^ exchanger 2 activity to reduce intracellular [H^+^] and improve Fz recruitment of Dsh^26^. Although serial replacement of basic DEP residues indicated that membrane stabilization of Dsh was more important to non-canonical PCP Wnt signaling than canonical Wnt/β-catenin signaling, it is reasonable that Dvl membrane stabilization by an alkaline pH_i_ in CF cells should also facilitate canonical Wnt/β-catenin signaling. Live imaging of enteroids from WT/ or Cftr KO/Dvl2 KO/Dvl2-EGFP/Rosa^mT/mG^ mice supported this hypothesis by showing greater juxtamembrane association of Dvl in Cftr KO CBC cells. Dvl membrane association was both charge- and pH_i_-dependent and could be predictably manipulated by pharmacological treatments to reduce intracellular [Cl^−^] in Cftr KO cells or inhibit Cftr in WT cells. Indeed, the acute change in Dvl2-EGFP membrane localization upon alkalization with Cftr_inh_-172 in WT enteroids is more consistent with an epithelial intrinsic effect of Cftr loss on increasing proliferation than genetic/epigenetic effects resulting from life-long absence of Cftr in the Cftr KO model.

The demonstration of greater Dvl association with the plasma membrane in Cftr KO CBCs in situ is consistent with the central hypothesis, but does not directly establish increased Fz-Dvl interaction. Quantitative information regarding the amplification of signal through the cascade of the Wnt/β-catenin pathway and discrete genetic manipulations of an appropriate cell line will be needed to estimate the contribution of Dvl membrane stabilization to increased Wnt/β-catenin signaling. A recent study investigating inflammation in the ΔF508 Cftr mouse small intestine concluded, in contradiction to our findings, that loss of Cftr activity suppressed active β-catenin signaling as compared to WT^83^. This apparent discrepancy may be methodological in that our measurements of active β-catenin expression were confined to lysates of freshly isolated crypts to evaluate the proliferative intestinal compartment. In contrast, the study of Liu et al. employed lysates of whole thickness small intestine in measurement of total and active β-catenin which would include the contribution of non-epithelial cell types.

An unexpected outcome in studies of WT CBCs was increased Dvl membrane association at the lateral-Lateral cell membrane. Although the cause of this phenomenon was not investigated, it is tempting to speculate that the lateral-Lateral cell membrane generates a localized pH_i_ gradient that increases Dvl localization. To the best of our knowledge, asymmetric cellular pH_i_ gradients in intestinal stem cells have not been reported. However, an asymmetric cellular pH_i_ gradient created by Na^+^/H^+^ exchanger 1 activity exists at the leading edge of migrating fibroblasts which is necessary for efficient activity of the Rho GTPase Cdc42^84, 85^. Cdc42 also plays a critical role in coordinating migration of mouse small intestinal stem cells^86^. Another potential function for asymmetric Dvl localization may also relate to Dvl’s role in mitotic spindle formation^87^, an orderly process that is associated with asymmetric division in normal intestinal stem cells but not under precancerous conditions, e.g., APC^min^ mice^88^.

Increased Dvl association with the supranuclear lateral plasma membrane in Cftr KO crypt epithelium may facilitate non-canonical Wnt pathways of planar cell polarity (PCP) and directional cell movement^89, 90^. Increased cell migration from the crypts as a consequence of increased epithelial proliferation has been shown in the Cftr KO mouse in vivo^14^. In intestinal crypts, cell migration must be closely coordinated with proliferation and involves remodeling of the adherens junctions by processes that are not well understood in this context^91^. Dvl is a crucial component of the PCP pathway as previously shown for polarized cell migration of mouse embryonic fibroblasts^92^. Dvl transduces signals from Wnt-liganded Fz and the co-receptor Ror2, a tyrosine kinase-like orphan receptor, which directs c-Jun N-terminal kinase (JNK) for activation of c-Jun, resulting in cytoskeletal reorganization and polarized cell migration^27^. Studies in Xenopus show that polarization of ectodermal cells involves xDsh accumulation with Fz7 at the apical adherens junctions in response to Wnt11^93^. Dvl may also provide a link between the Fz/Ror2 pathway and Cdc42 for polarized cell migration. The Rho family GTPase Cdc42 is a major determinant of apical-basal polarity in intestinal epithelium^86^. Cdc42 is recruited by guanine exchange factors, a process facilitated by an alkaline pH_i_,85 and can bind partitioning complex members Par3 and Par6 at the tight junctional complex and apical membrane, respectively^94^. This process leads to activation of an atypical protein kinase C (aPKC, PKCζ), potentially regulated by the DEP domain of Dvl^27^, which facilitates polarized cell migration^95^. Additional studies will be necessary to sort out the cellular processes that coordinate crypt cell proliferation with crypt cell migration.

In conclusion, our data demonstrate that the absence of Cftr activity in murine ISCs establishes an alkaline pH_i_ that potentially facilitates canonical Wnt/β-catenin signaling by increasing the stability of the Wnt transducer Dvl at the plasma membrane. The rate of ISC proliferation and changes of signaling in the Cftr KO intestine are moderate, but over the course of a lifetime create an insidious process which is predicted to raise the risk of intestinal carcinogenesis. When placed within the context of the Cftr KO intestinal environment, with attendant inflammation and dysbiosis, a significant risk for neoplasia develops which is consistent with the high tumor penetrance (~60%) exhibited by aged CF mice^13^. However, to best of our knowledge, epithelial proliferation rates and Wnt/β-catenin signaling of the intestine have not been evaluated in CF patients. These data from the Cftr KO mouse model should illuminate the need to investigate this potential risk factor in CF. With continued improvements in care and an aging CF population, the incidence of digestive tract cancer will likely increase. There is a need for more attention to clinical screening of CF patients for GI cancer and investigations identifying the underlying processes by which life-long dysfunction of an epithelial anion channel yields increased gastrointestinal cancer risk.

## Acknowledgements

The authors would like to acknowledge the assistance of the Dalton Cardiovascular Research Center Live Cell Imaging Core and Dr. Luis Martinez-Lemus (director).

